# Invariant Genes in Human Genomes

**DOI:** 10.1101/739706

**Authors:** Ankit Kumar Pathak, Ashwin Kumar Jainarayanan, Samir Kumar Brahmachari

## Abstract

With large-scale human genome and exome sequencing, a lot of focus has gone in studying variations present in genomes and their associations to various diseases. Since major emphasis has been put on their variations, less focus has been given to invariant genes in the population. Here we present 60,706 genomes from the ExAC database to identify population specific invariant genes. Out of 1,336 total genes drawn from various population specific invariant genes, 423 were identified to be mostly (allele frequency less than 0.001) invariant across different populations. 46 of these invariant genes showed absolute invariance in all populations. Most of these common invariant genes have homologs in primates, rodents and placental mammals while 8 of them were unique to human genome and 3 genes still had unknown functions. Surprisingly, a majority were found to be X-linked and around 50% of these genes were not expressed in any tissues. The functional analysis showed that the invariant genes are not only involved in fundamental functions like transcription and translation but also in various developmental processes. The variations in many of these invariant genes were found to be associated with cancer, developmental diseases and dominant genetic disorders.

## INTRODUCTION

The international efforts to sequence the human genome have thrown up some surprising findings, that will take a long time in order to completely understand the human genome and its function (1, 2). With the advent of large scale whole genome and exome sequencing projects in the past decade, we have seen an unprecedented scale of variation in the protein-coding genes across the human population (3). The variations that exist in the human genome are the keys in determining the phenotype of an individual. Such variations can be a difference in a single nucleotide (single nucleotide polymorphism) or in thousands of bases (structural variation) (4). SNPs can have a profound effect on gene expression, the amino acid sequence and consequently phenotype (5). Structural variations that include deletions, insertions, duplications and translocations are also responsible for generating diversity within the human population (6). The genetic variations present in the human genome are responsible for distinctive characteristics of an individual and thus these variations have made it unrealistic to define a common human genome.

Although much effort has gone into deciphering the relationship between genomic variations and phenotype, very little effort has been put in understanding what are the true invariant genes in the large human genome sequence datasets, both population specific and across the populations. Identification of highly conserved non-coding regions has led to the discovery of miRNA and other conserved non-coding RNA that have been shown to have major regulatory functions in the biological processes (7–11). In order to address invariant genes in human genomes across the populations, our work focuses on protein-coding genes with extremely low (allele frequency less than 0.0001) variation rates in large population specific datasets termed as ‘invariant’ coding regions. These genes have synonymous variations but lack any amino acid variation in exons. Studying gene statistics, functional classes, expression patterns and homology of these invariant genes in the human genome will help to reveal the conserved protein functions across human populations. We have identified 423 invariant proteins in 60,706 human genome spread over 6 subpopulations with varied presence in different populations. To our surprise, of the 46 absolute invariant proteins that we identified, 8 of the proteins are unique to human genome wherein 3 of them still have unknown functions. Further, looking at the invariance patterns in human subpopulations has revealed specific invariant proteins unique to subpopulations arising out of natural selection. Although we don’t have any cancer genome representation in the dataset, we notice these subpopulation specific invariant genes to be associated with several cancers and dominant genetic disorders.

## MATERIALS AND METHODS

### Databases

#### Exome Aggregation Consortium (ExAC)

The exome sequencing data was retrieved from Exome Aggregation Consortium (ExAC) database (Release 1.0) (12) with 10,195,872 variations in 60,706 unrelated individuals aggregated from various population genetic studies. The dataset was inclusive of samples from individuals with adult-onset diseases such as Type 2 Diabetes and Schizophrenia while no samples with tumors or severe pediatric diseases were accounted in the dataset. The database provided population-wise allele frequencies (AF) of all variations for African/African Americans (5,203 individuals), Latinos (5,789 individuals), East Asians (4,327 individuals), Finnish (3,307 individuals), Non-Finnish Europeans (33,370 individuals), South Asians (8,256 individuals) and Others (454 individuals) subpopulations. The database uses GRCh37/hg19 reference human genome assembly.

#### RefGene

RefGene database from the UCSC Genome Browser Database (13) provided the reference coordinates and genomic sequences of known human protein-coding and non-protein-coding genes. It was used to analyze statistical features of exons for varied gene sets. The RefGene database uses BLAT-generated annotations to align multiple regions.

#### Protein ANalysis THrough Evolutionary Relationships (PANTHER)

PANTHER Classification System (Version 14.1) (14) was used for ontology-based functional classification and statistical overrepresentation of Invariant Genes. For each subpopulation, molecular functions and biological processes of the genes were assessed using the database. Fisher’s Exact test with False Discovery Rate (FDR) correction was used for statistical overrepresentation analysis of the genes.

#### Genotype-Tissue Expression (GTEx)

The Genotype-Tissue Expression (Version 7) database (15) provided tissue-specific gene expression data for 53 non-diseased tissue sites across nearly 1000 individuals. Median TPM (Transcripts Per Kilobase Million) values were used for all gene expression representations.

### Gene-based functional annotation of variants

Exonic variations provided by the ExAC database were passed through ANNOVAR (16), a tool to functionally annotate genetic variants. ANNOVAR, using RefGene database from the UCSC Genome Browser, mapped the variations to their corresponding genes and exons and further classified them into different classes of mutations depending upon their effect on proteins (Non-synonymous, Synonymous, Frameshift Deletion, Non-frameshift Deletion, Frameshift Insertion, Non-frameshift Insertion, Frameshift Substitution, Stoploss and Stopgain).

### Screening for Invariant Genes

To avoid low-quality variations in our analysis, a genotype quality score (GQ) >/= 20 and read depth score (DP) >/= 10 were used, as provided by the ExAC dataset. Out of 10,195,872 exome sequence variations, 10,192,197 variations passing the quality scores were used for all future analyses. Further, we removed all the personal mutations from the population specific dataset which had an allele frequency (AF) of less than equal to 0.0001. All genes were screened for any protein-altering mutation within the passing variations and the genes which carried only synonymous variations were taken up for subsequent analysis in a given subpopulation. From the above dataset of all subpopulations we eliminated all the genes with allele frequency variation of 0.001 and above to obtain the invariant genes across all subpopulations.

## RESULTS

### ExAC dataset

About 10 million protein-coding variations were provided by ExAC dataset with varied representation of human genome across African, Latino, Finnish, non-Finnish European, South Asian and East Asian geographic ancestries. The allele frequency distribution of the sequence variations revealed the abundance of personal mutations in the human exome where non-synonymous variations were present in substantially larger scale (Fig. 1). The average allele frequency histogram showcases the extent of non-uniform distribution of different functional classes of variations in the human population (Supplementary Fig. 1). Interestingly, the non-frameshift deletions/insertions mutations have a much higher allele frequency than non-synonymous variations.

**Figure 1.**
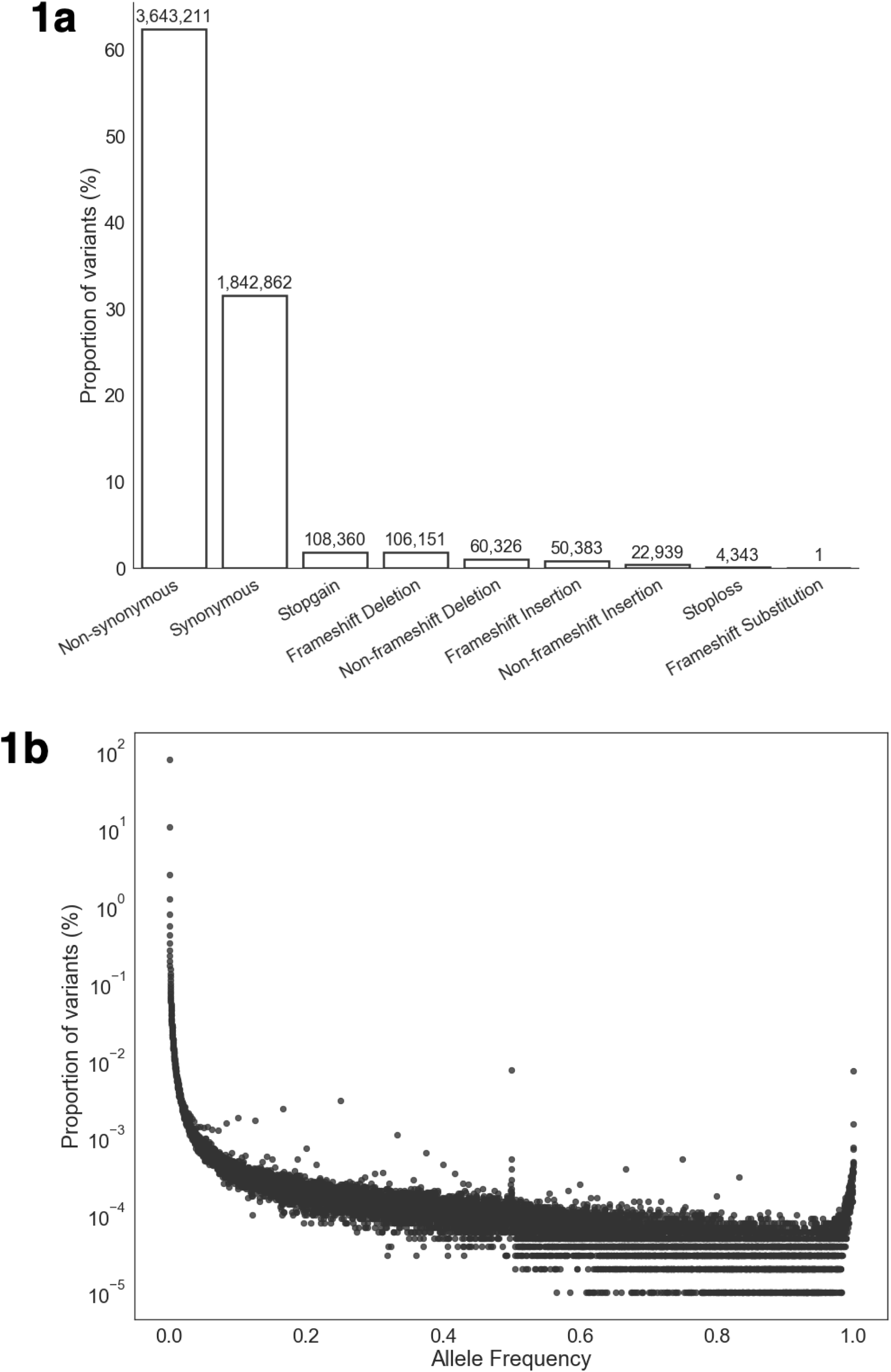
Variant statistics in the ExAC dataset. **a,** Number of mutations in the ExAC dataset classified by their impact on protein sequence. **b,** Distribution of all variations in the ExAC dataset with respect to their allele frequencies rounded off to 4 decimal places.

A subpopulation-wise allele frequency cutoff of 0.0001 and below was used to eliminate all personal mutations in the subpopulation. This dataset was used to identify invariant genes in subpopulations with no mutations leading to an amino acid change.

### Invariant genes

The analysis revealed genes that were invariant among different subpopulations and invariant genes unique to specific subpopulations as shown in Table 1. FIN, an isolated inbred subpopulation, had significantly higher number of these invariant genes than the other subpopulations with a broader gene pool. AFR, SAS and EAS had smaller invariant gene sets reflecting upon the diversity in the subpopulations. There was no correlation between the number of invariant genes and the number of individuals in the subpopulation samples (r = -0.03). These accounted for a total of 1,336 genes which were invariant in at least one subpopulation.

**Table 1.**
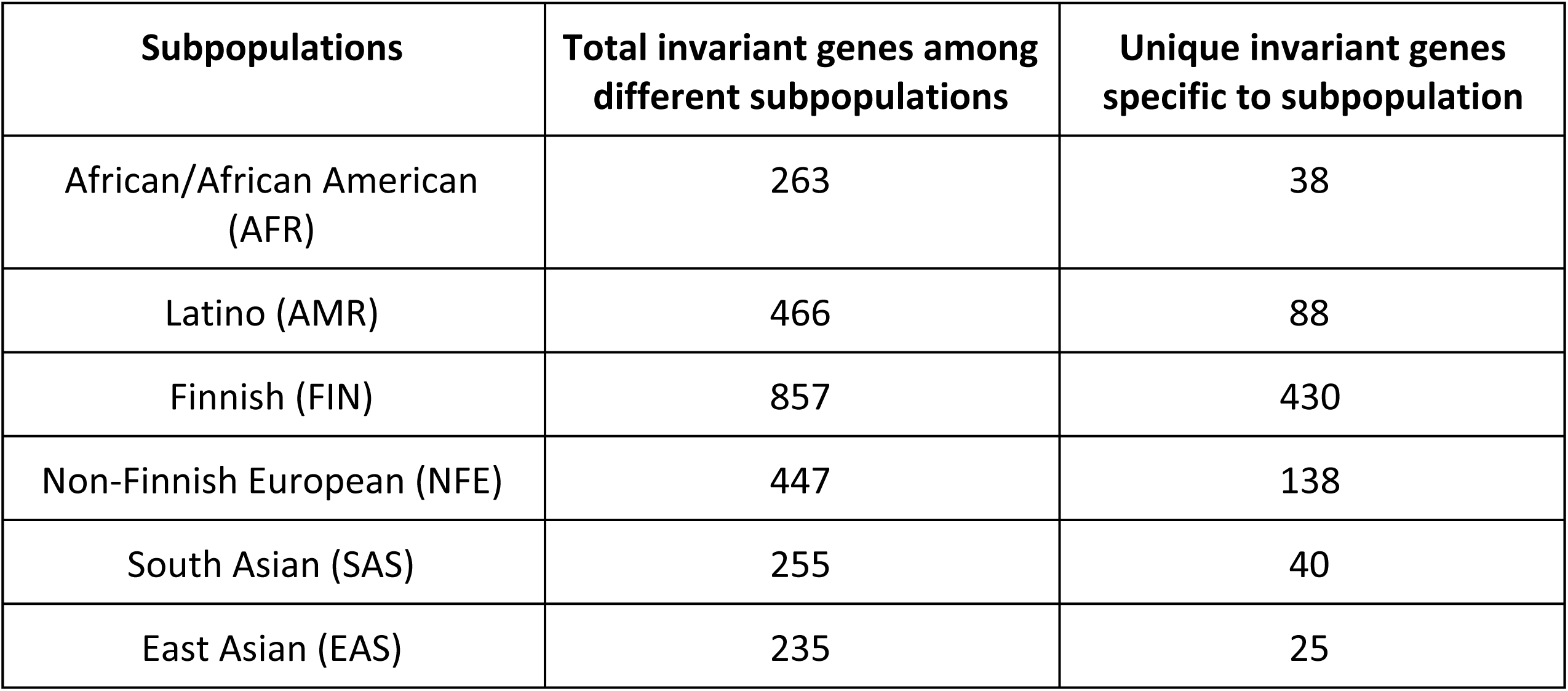
Invariant genes across subpopulations. A union set of 1,336 genes whose invariance across different subpopulations is shown. A detailed list of invariant genes is in Supplementary File 1.

Out of the 1,336 total genes, drawn from different subpopulations, some of the genes exhibited low frequency/common (allele frequency above 0.01) variations in other subpopulations (17). When a cutoff of 0.01 and above was applied to these set of genes, a total of 878 genes were obtained (Supplementary File 2). However, we wanted to apply a more stringent cutoff frequency (0.001 and above) to obtain 423 invariant genes, even though this resulted in removal of some of the rare variants (17). These 423 invariant genes accounting to 2094 exons had allele frequencies within the range of 0.0001 and 0.001 for all subpopulations (Supplementary File 3). 46 invariant genes had no observed mutations in the entire population studied.

With a mean of ∼5 exons, all subpopulations showed a similar distribution for the number of exons per invariant gene (Supplementary Fig. 2) which is much less than a global average of ∼9.5 exons per gene in humans (18). The length of exons in invariant genes were also similarly spread in subpopulations with an average ∼420 base pairs per exon (Supplementary Fig. 3) as compared to a global average of ∼315 base pairs per exon for all human genes.

#### Functional Classification

Biological processes ontology of the invariant genes among different subpopulations showed their involvement majorly (> 80%) in cellular process, metabolic process, biological regulation and localization. Other classes like multicellular organismal process, developmental process, reproduction, response to stimulus, cellular component organization and biological adhesion were also observed to be present in all subpopulations (Fig. 2a). While binding and catalytic activity predominantly encompassed the molecular functions, other activities like molecular function regulator, molecular transducer activity, structural molecule activity, translation regulator activity, transcription regulator activity and transporter activity were also observed in all subpopulations (Fig. 2b). The ontologies profile of the 423 invariant genes also exhibited similar classes of biological processes and molecular functions (Fig. 3a, 3b).

**Figure 2.**
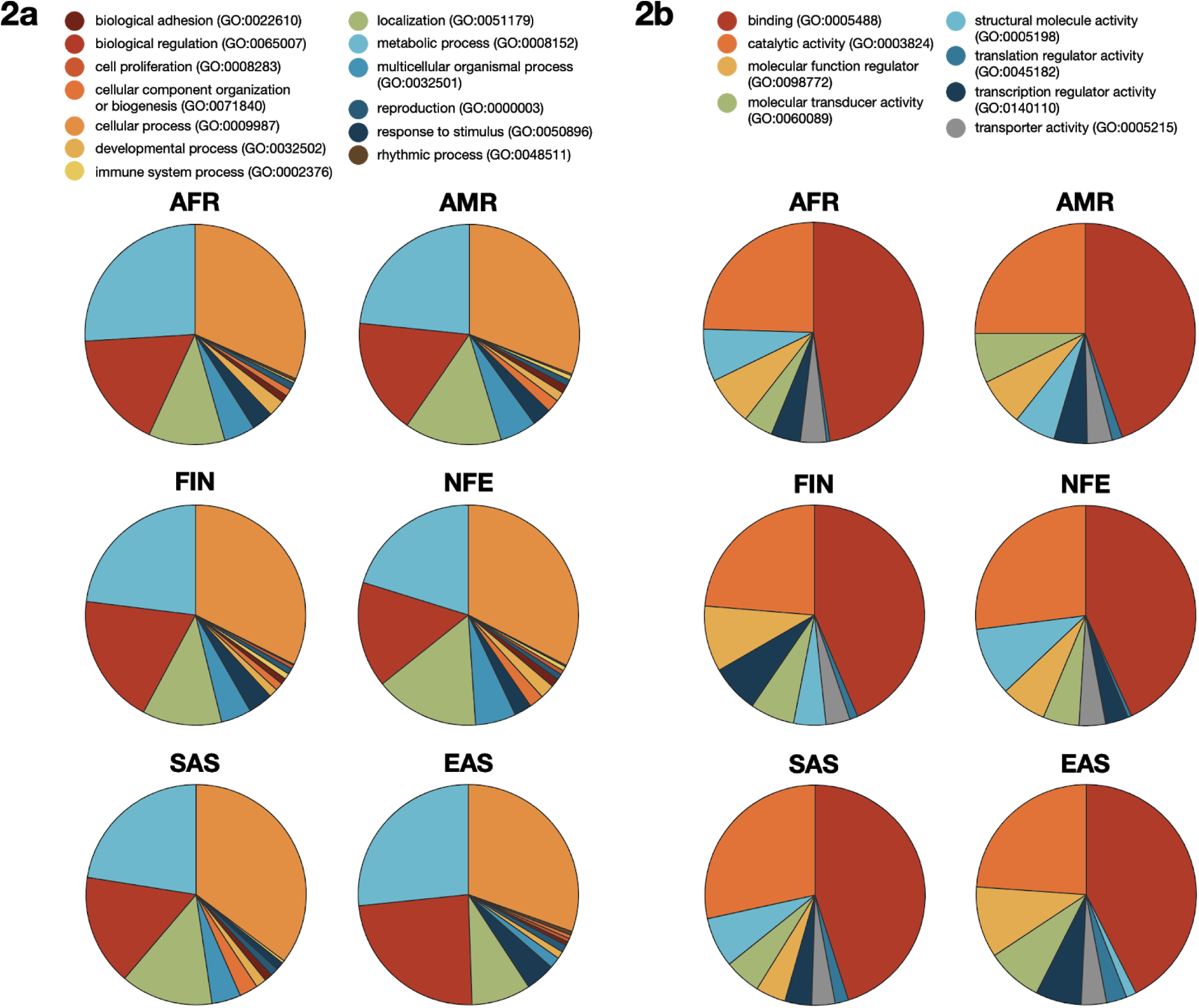
Ontological functional classification of invariant genes across subpopulations. **a,** Share of biological processes performed by invariant genes across subpopulations. **b,** Share of molecular functions performed by invariant genes across subpopulations.

**Figure 3.**
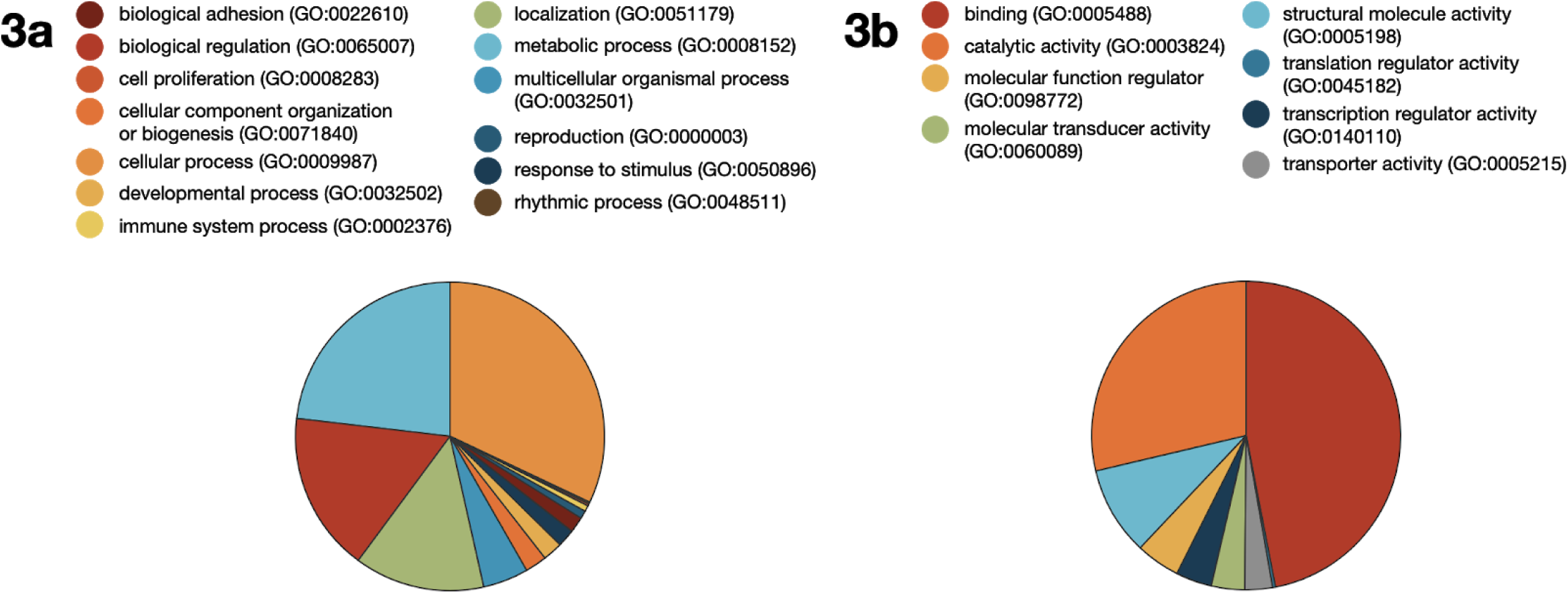
Ontological functional classification of 423 invariant genes. **a,** Share of biological processes performed by invariant genes. **b,** Share of molecular functions performed by invariant genes.

Overrepresentation tests showed pathways in core biological processes like splicing, proteolysis, apoptosis, chromatin/chromosome organization and localization highly enriched in the invariant genes in the subpopulations. Pathways associated with Huntington and Parkinson disease were also depicted to have a heightened involvement while integrin signalling, CCKR signaling, FGF signaling and cytoskeletal regulation by Rho GTPase pathways were also featured. Molecular activities that were enriched were associated with transcription and translation initiation factors, ubiquitin protein ligases, structural constituent of ribosomes, phosphatase inhibitors and receptor inhibitors (Supplementary File 4a-c).

#### Gene Expression

Expressions profiles of invariant genes among different subpopulations retrieved from GTEx Portal revealed their preferred expression across all tissues. About 69% in AFR, 73% in AMR, 66% in FIN, 79% in NFE, 74% in SAS and 50% in EAS of invariant genes were ubiquitously expressed in the provided 53 non-diseased tissues (Supplementary File 5). Out of the 423 invariant genes, only 403 showed GTEx expression profiles of which 329 genes (81%) showed expression across all tissues whereas 31 invariant genes showed no expression across any tissue (Supplementary File 6).

### Absolutely invariant genes common to all subpopulations

The list of 46 absolutely invariant genes common to the 6 subpopulations is shown in Table 2. The X chromosome had significantly higher share of these genes than the autosomes. The length of the proteins ranged from 1,138 amino acid residues and skewed towards the lower values with the smallest protein of just 51 amino acid residues. Uncharacterized genes such as LOC101928120, LOC730098 and C18orf12 were also seen to be absolutely invariant in all subpopulations but no annotated protein families and domains were found. Multiple copies of some of these genes were found to be clustered - CT47A family with 10 genes, USP17L family with 9 genes, GAGE family and PPIAL4 family with 3 genes, SPACA family with 2 genes. Most of these genes were traceable to species closely related to humans, however 8 genes were unique to the human genome.

**Table 2.**
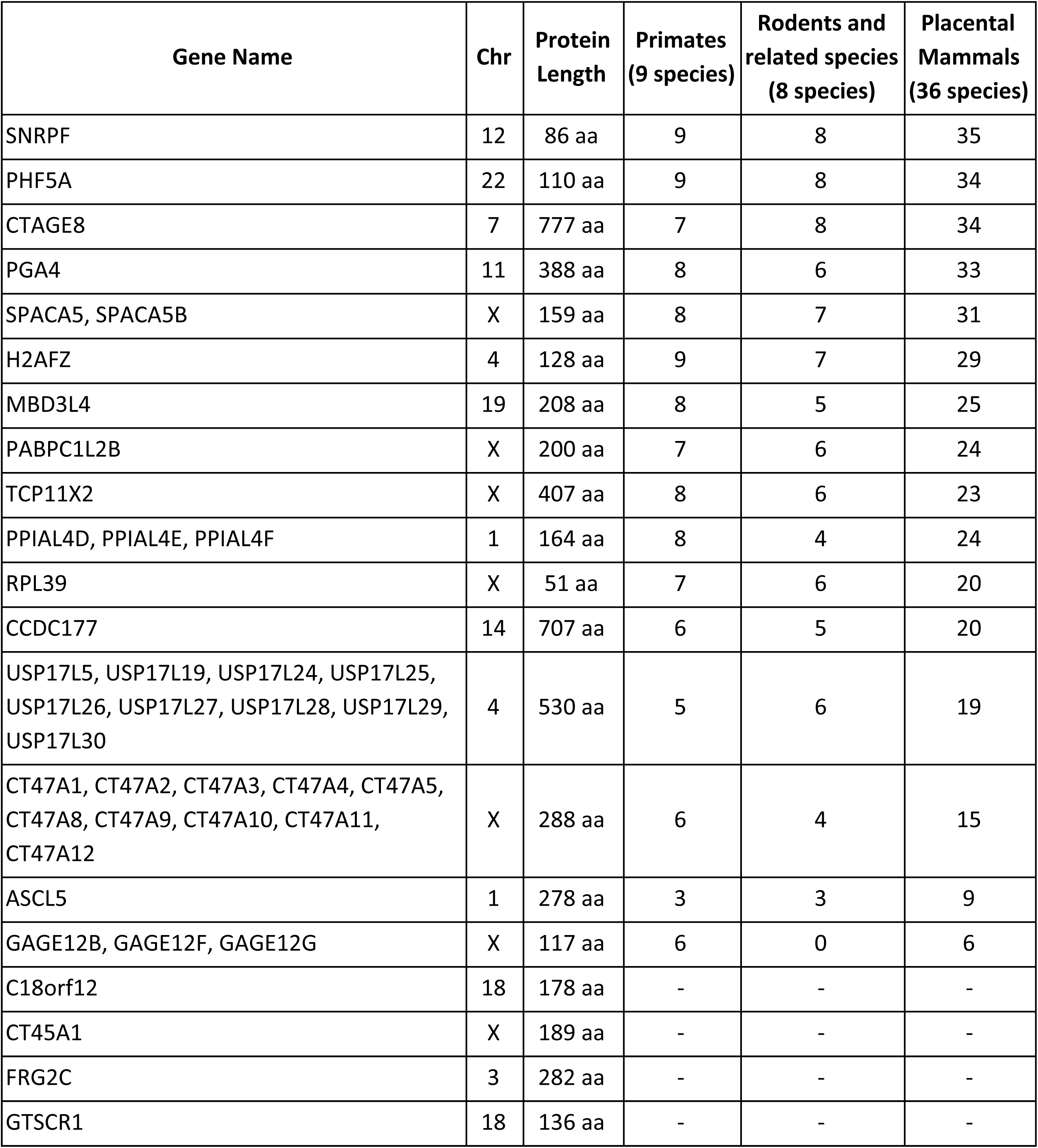
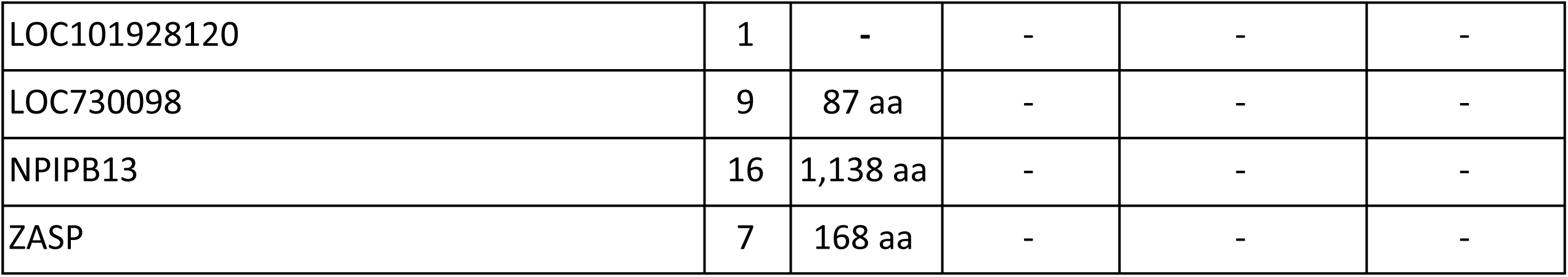
Absolute invariant genes common to subpopulations.

#### Functional Classification

The 46 absolutely invariant genes had conserved biological processes in components of biological regulation (regulation of apoptotic process), cellular process, reproduction (spermatogenesis), cellular component organization/biogenesis, localization, metabolic process and response to stimulus (defense response to bacterium). Molecular functions in these genes were observed to have binding, catalytic activity, transcription regulator activity and transporter activity to be conserved. (Fig. 4a, 4b)

**Figure 4.**
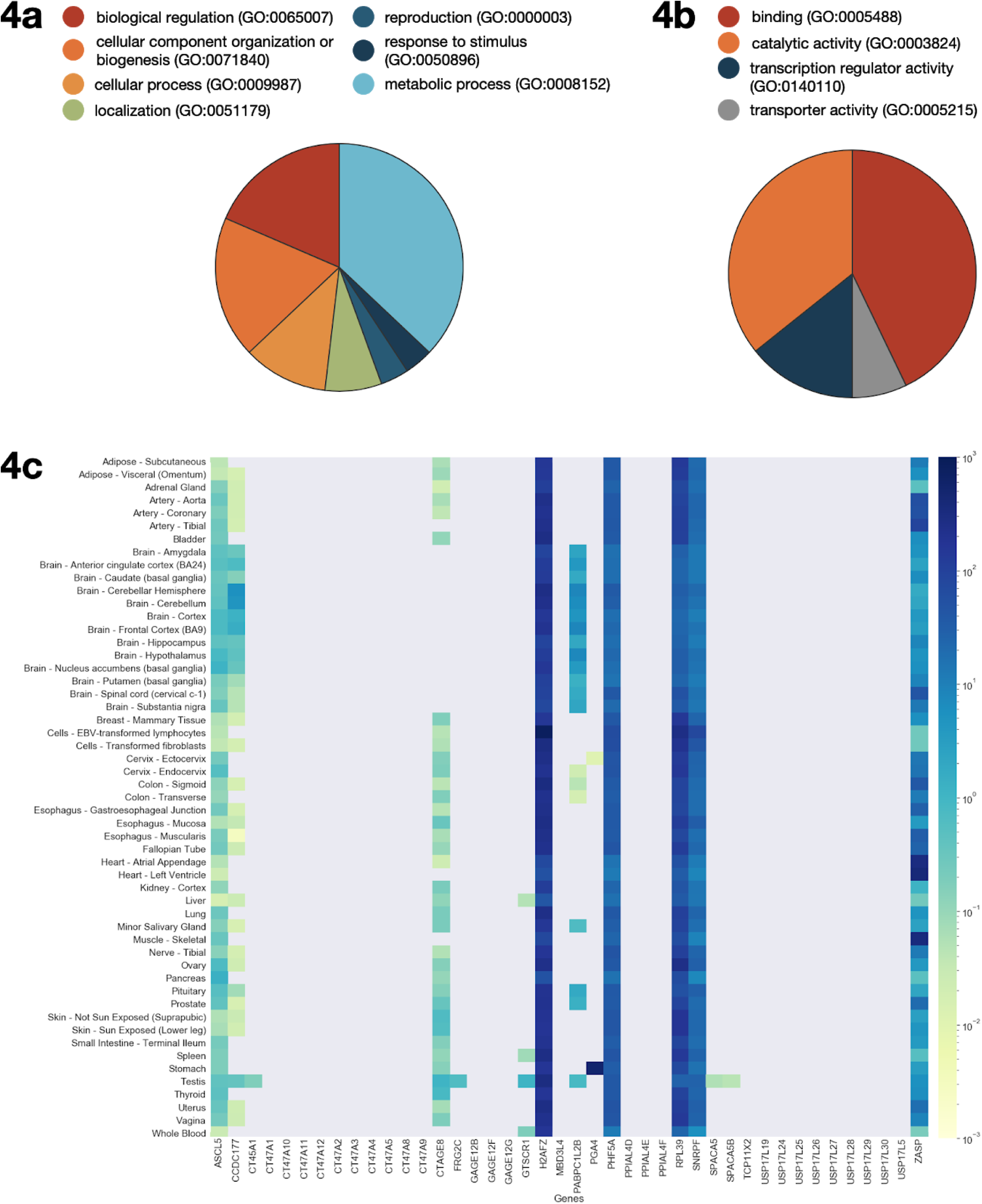
**a**, Share of biological processes performed by absolute invariant genes common to all subpopulations. **b**, Share of molecular functions performed by absolute invariant genes common to all subpopulations. **c**, Gene expression profiles across 53 non-diseased tissues in log-scaled Transcripts Per Million (TPM) values ranging from 0.01 to 550 for absolute invariant genes common to all subpopulations.

#### Gene Expression

The expression levels of the absolutely invariant genes can be broadly classified into three groups; genes that are ubiquitously expressed in all tissues (ASCL5, H2AFZ, PHF5A, RPL39, SNRPF, ZASP), genes that are expressed across various tissues (CCDC177, CTAGE8, GTSCR1, PABPC1L2B), genes that are tissue-specifically expressed (CT45A1, FRG2C, PGA4, SPACA5/SPACA5B). Surprisingly, CT45A, USP17L, GAGE and PPIAL4 gene families did not show any expression in the GTEx Portal or in mouse embryonic expression dataset (19). The gene expression profiles for C18orf12, LOC101928120, LOC730098 and NPIPB13 were not available in the GTEx Portal (Fig. 4c).

#### Functional analysis of unique invariant genes

Among the 46 absolutely invariant genes, essentiality of the few have been addressed in detail below.

##### PHD finger-like domain-containing protein 5A (PHF5A)

Embryonic stem cells (ESCs) are pluripotent and capable of giving rise to all tissues of the embryo through the process of differentiation. This process is governed and regulated by cell-specific factors resulting in differential gene expression. One such factor is PHF5A, a highly conserved protein containing a plant homeodomain (PHD) domain. A recent study shows that PHF5A protein level depletion results in loss of pluripotency in ESCs and inhibition of reprogramming (20). Another study shows that knockdown of PHF5A gene in C.elegans results in abnormal organogenesis during development (21). Therefore, it is highly probable that the multitasking role of PHF5A in important processes during development.

##### Achaete-scute (ASCL)

ASCL (achaete-scute complex-like) is a gene family that includes five members (ASCL1, ASCL2, ASCL3, ASCL4 and ASCL5). All the members of this gene family encode transcription factors that control nervous system development (22). Studies also show that they are involved in cell fate determination of neuroblast (23). Yet, the role of ASCL members does not remain confined to nervous system development. They have also been found to be expressed in progenitor cells during the differentiation of muscle and gut and this shows the significance of ASCL expression during organogenesis (24). It can thus be said that the role of ASCL in several important developmental processes has made it highly conserved.

##### ZO-2 associated speckle protein (ZASP)

ZASP is a protein containing PDZ-LIM domain and is expressed mostly in the striated muscle. This domain is responsible for protein-protein interactions which enable the protein to form multiprotein complexes and ZASP is required to maintain the structural integrity of sarcomeres during contraction (25). Since mutations in this protein result in myopathies and because ZASP plays an important role in sarcomere, it remains essential.

##### Histone H2A.Z (H2AFZ)

H2AFZ is a variant of histone H2A and mediates thermosensory response. Usually, variant histones replace canonical histones under certain circumstances and result in a change in chromatin structure and function which in turn alters gene expression. A recent study shows that H2AFZ acts as a recruitment platform for many proteins that play a role in gene regulation and its binding to PWWP2A specifically regulates mitosis and neural crest differentiation (26). Thus, the crucial role of H2AFZ in regulation of gene expression has made it essential in humans.

##### Ubiquitin Specific Peptidase 17 (USP17)

USP17/DUB3 is a deubiquitinating enzyme which cleaves ubiquitin from precursor protein. Studies have shown the importance of this enzyme in cell growth and survival (27). Also, another study shows the requirement of USP17 in virus-induced type I IFN signaling (28). Because of its regulation in important processes, the lack of variations in USP17 gene is understandable.

Some of the oncogenes in humans includes *CT45A1, CT47, CTAGE8, GAGE* and *RPL39*. Studies have shown that mutations in such oncogenes affects cell division and proliferation (29–34), thereby making these genes so significant in humans.

### Unique invariant genes specific to each subpopulation

An inbred FIN subpopulation had 430 genes seen to be specifically invariant among its individuals whereas the diverse EAS subpopulation accounted for the least number of 25 exclusively invariant genes. We further investigated the 430 invariant genes specific to isolated inbred FIN subpopulation for their low-frequency/common variants (allele frequency greater than 0.01) across the other subpopulations (Supplementary File 7). The analysis revealed that 259 out of 430 invariant genes specific to FIN varied in at least one subpopulation. 73 of these unique invariant genes showed variation in all subpopulations while 109 showed variation in only one subpopulation. The variation seen in these genes is due to the environmental selection acting in the respective subpopulation. These genes can be used to understand the role of environmental factors and natural selections in other subpopulations.

#### Functional Classification

Components of biological processes including biological regulation, cellular component organization, biogenesis, developmental process (anatomical structure development), localization, metabolic process and response to stimulus (response to stress) were observed in each subpopulation. Biological adhesion (cell adhesion) was observed in all subpopulations except AFR whereas cellular processes including cell cycle process, cellular metabolic process, signal transduction were absent from EAS while present in the rest. Cell proliferation and locomotion (cell motility, taxis) were seen in AMR, FIN and NFE and multi-organism process (response to other organism) was present in AMR and NFE. Conserved processes in immune system, multicellular organismal process (multicellular organism development, system process, pattern specification process), reproduction (gamete generation) and signaling (cell-cell signaling, signal transduction) were observed in AMR, FIN, NFE and SAS but absent in AFR and EAS (Supplementary Fig. 4).

Subpopulation specific invariant genes were also involved in several classes of molecular functions, predominantly binding activity and catalytic activity. Heat shock protein binding was solely observed in AFR. Oxidoreductase catalytic activity was observed only in AMR, FIN and NFE. Molecular function regulator activity showed variation in AFR, AMR, NFE and SAS while structural molecule activity varied in AFR, AMR, FIN and EAS subpopulations in much greater extent. Molecular transducer activity entailing neurotransmitter receptor activity was profoundly active in AMR, FIN and NFE while was not observed in EAS. Transcription regulator activity was also varied in EAS while largely conserved in FIN subpopulation. Also, transporter activity was found to be varied in SAS. Regulation at the level of translation was also conserved to a lesser extent in the FIN subpopulation (Supplementary Fig. 5).

The essentiality of a few unique invariant genes specifically belonging to the 6 subpopulations have been addressed in detail in Table 3.

**Table 3.** Essentiality of invariant genes belonging specifically to different subpopulations.

| Gene Name | Population | Description proving essentiality |
| --- | --- | --- |
| CABP1 | AFR | Malaria is one of the most prevalent diseases in Africa. Studies have reported almost 50% of malarial patients exhibiting neurological deficits in sub-Saharan Africa. Recent studies have revealed CABP1, a neuron-specific regulator of calcium channel activation, to exhibit an increased gene expression in malaria infected brains resulting in neurological diseases. (35,36) |
| ELOVL6 | AFR | Triple-negative breast cancer is more aggressive than other subtypes of breast cancers and has a lower survival rate. It has the highest incidence in African American women. A recent study found mRNA expression levels of ELOVL6, a fatty acid elongase, to be significantly higher in these tumors. (37) |
| GNPDA2 | AFR | African-American women have the highest obesity rates as compared to other groups in the United States. Studies exploring body mass index (BMI) related loci in Africans revealed GNPDA2 to be significantly associated with an increase in BMI (38–40). This gene has a single mutation (R23C) with an allele frequency of 0.03 in FIN subpopulation (Supplementary File 2). |
| PAFAH1B3 | AFR | Native Ethiopians residing at 3,500 meters above sea level on Bale Plateau or Chennek field in Ethiopia need high altitude adaptations. In recent studies, PAFAH1B3 has been shown to modulate hypoxia tolerance amongst Ethiopian highlanders. (41) |
| TNFSF4 | AFR | Systemic lupus erythematosus (SLE) is a systemic autoimmune disease which causes autoantibody production, abnormalities of immune system function, and damage in several organs. SLE is 4 times more common in individuals of African-American ancestry than those of European ancestry. In recent reports, mutations in TNFSF4 have been associated with SLE amongst African populations (42,43). This gene has a single mutation (V27M) with an allele frequency of 0.01 in SAS subpopulation (Supplementary File 2). |
| ARL4C | AMR | Males over the age of 65 and tobacco using population in the United States are more susceptible to abdominal aortic aneurysm (AAA), a dilatation of the infrarenal aorta. It is noted that the differentially expressed gene ARL4C expression is upregulated in AAA of American type culture versus controls. This gene was not previously implicated in AAA pathobiology. This gene could probably have a significant role as the treatment and diagnostics tool for AAA and hence, this gene is specific to Americans as they are more prone to this disease. (44) |
| CMTM7 | AMR | From the Global cancer statistics, the risk of being diagnosed with cancer, especially lung and breast cancer is highest in America especially the North America. Recent findings suggest that CMTM7 is a novel 3p22 tumor suppressor regulating G1/S transition and EGFR/AKT signaling during tumor pathogenesis. CMTM7 is associated with the inhibition of EGFR-PI3K/Akt signaling pathway, resulting in the upregulation of p27 and further inhibition of CDK2 and CDK6. CMTM7 may inhibit cell proliferation and migration via the EGFR-PI3K/Akt pathway. Thus, CMTM7 which is a potent TSG and has important suppressive role in cancer development and progression must be an essential gene among Americans. (45) |
| COLCA2 | AMR | The colorectal cancer is one of the common types of cancer. COLCA2 is present in the cytoplasm of normal epithelial, immune and other cell lineages, as well as tumor cells but its expression is reduced in tumor cells from subjects with higher risk alleles. It is possible that COLCA2 has critical functions that suppress tumor formation in epithelial cells. Elevated levels of COLCA2 in immune and other cells of the microenvironment may also provide protection against cancer cell growth. In a recent study it is implicated that COLCA2 has a major role in the pathogenesis of colon cancer and also in the involvement of immune pathways. Functional validation of susceptibility locus 11q23 correlated with the expression of the gene COLCA2 by the immune system in the samples from African Americans. We also observe an increase in COLCA2 protein expression in colon tissues from individuals with the |
|  |  | lower risk African Americans genotype. Thus, the specific expression of COLCA2 gene in African Americans serves as a tool against colorectal cancer (46–48). This gene has a single mutation (Y174X) with an allele frequency of 0.03 and 0.02 in AFR and NFE subpopulations respectively (Supplementary File 2). |
| COX5A | AMR | The recent study showed that nondiabetic Northern European Americans were substantially more insulin sensitive than both diabetic and nondiabetic Indians. The gene COX5A involved in oxidative phosphorylation pathway was noted to be upregulated in Asian Indians compared with European Americans. This gene may probably account for the less insulin resistant aspect of European Americans compared to that of Asian Indians. It is also commented that COX5A gene is differentially methylated between African American diabetes patients without nephropathy and end stage renal disease patients being treated by hemodialysis. Additionally, the differentially expressed gene COX5A was observed to be upregulated in African American patients with prostate cancer. Thus, these reports suggest that COX5A gene is specific among American population. (49–51) |
| DAD1 | AMR | Asthma is classified as a complex, chronic and inflammatory disease of the respiratory tract. Several genome-wide association studies have been carried out in recent years and it was noted that the 17q21 region is associated with asthma symptoms in childhood, but the study on Latin American population expressed a different result where 14q11 and 15q22 regions were found to be associated and 17q21 region was not found to be associated with asthma symptoms in childhood. The gene DAD1 which is present in 14q11 region is found to be active in apoptosis regulation process and its failure in this process may lead to increased lymphocytes in asthma patients. Thus, this gene DAD1 is most likely found specific among Latin American population. (52) |
| DOLPP1 | AMR | Nephrolithiasis is a common problem worldwide with an increasing incidence in Westernized societies. The prevalence of kidney stones was figured to be increased in American adults and this may be due to environmental factors. Recurrent calcium stone-forming patients have been shown to excrete increased quantities of abnormal Tamm-Horsfall protein (THP), with a change in its chemical composition to include more sialic acid residues. Maturation of THP may be regulated by other genes like DOLPP1 and this again represent a candidate gene for nephrolithiasis more specifically among US population (53,54). This gene has a single mutation (A178T) with an allele frequency of 0.03 in FIN subpopulation (Supplementary File 2). |
| HRH2 | AMR | After observing different races, certain genotypes were overrepresented among children with allergic asthma. Among them HRH1-17 (HRH2 543 G/A) genotype was observed to be overrepresented among African-American children with allergic asthma. Thus, HRH2 gene may be of particular importance in the pathogenesis of African American population with allergic asthma. It is also demonstrated that HRH2 gene is differentially methylated between African American diabetes patients without nephropathy and end stage renal disease patients being treated by hemodialysis (50,55). This gene has a single mutation (N266S) with an allele |
|  |  | frequency of 0.04 in FIN subpopulation (Supplementary File 2). |
| EIF4EBP1 | FIN | Type II Diabetes is one of the complex diseases, where most disease-associated variants identified by Genome-wide association studies are different across different populations. In this study it was figured out that the gene product of EIF4EBP1 is involved in protein-protein interaction network associated with Type II Diabetes specifically among Finnish population. (56) |
| GDAP1 | FIN | Charcot-Marie-Tooth (CMT) disease is the most common hereditary neuromuscular disorder. Earlier it was found that mutation in GDAP1 cause axonal CMT disease by disturbing mitochondrial fission and during the study of prevalence of CMT disease in a population-based sample in Finland it was noticed that the founder mutation p.His123Arg in the GDAP1 gene to be the most common etiology for CMT2. Thus, we recognise the specificity of GDAP1 gene in Finnish population. (57) |
| POU3F1 | FIN | Multiple sclerosis (MS) is the most common autoimmune demyelinating disease of the central nervous system. In a study the genes responsible for MS by studying monozygotic (MZ) twin pairs discordant for MS was identified and over two-fold up-regulation of six genes in 40% of MZ MS Finnish twins suggests their role in MS pathogenesis and one among them is POU3F1. The protein (SCIP/Oct-6) translated by this gene has been found majorly in the nervous system, where it is associated with Schwann cells in the process of remyelination. Furthermore, it was revealed that misfunction of this gene leads to persistent hypomyelination and eventual axonal loss but no evidence of demyelination/remyelination processes or impaired Schwann cell proliferation was revealed. (58,59) |
| NDUFA4 | FIN | NDUFA4 was found among the other differentially expressed genes in Multiple sclerosis dataset in Finnish cohort. NDUFA4 has many protein-protein interactions including PARK7. These differentially expressed genes were also concluded to regulate epigenetic and apoptotic pathways that may further elucidate underlying mechanisms of autoreactivity in Multiple sclerosis. (60) |
| MTRNR2 | FIN | Mutations in MTRNR2 gene in mitochondrial DNA can cause hearing impairment. The most prevalent one is m.1555A > G in this gene. In a study the frequency of this mutation among the Finnish adult patients with matrilineal sensorineural hearing impairment was found to be 2.6%. Population studies suggests that this is common among the Finnish population. (61) |
| MSX1 | FIN | Hypodontia, congenital lack of one or a few teeth, is an autosomal dominant trait. On the other hand, the term oligodontia commonly refers to the congenital absence of more than six teeth. The study in Finnish population reported that MSX1 and MSX2 play important roles in the initiation and early morphogenesis of teeth. Although a defect in MSX1 is not involved in incisor and premolar hypodontia, it was conceivable that a mutation of the gene may cause other types of hypodontia in humans. In another study on Finnish population it was found that MSX1 mutations do not cause oligodontia. But in humans, in addition to anodontia, it is suggested that a mutation in MSX1 may be involved in cases where hypodontia is associated with cleft palate or other craniofacial malformations (62,63). This gene has a single mutation (A40G) with an allele frequency of 0.15 in AFR, 0.20 in AMR, 0.20 in NFE, |
|  |  | 0.13 in SAS and 0.03 in EAS subpopulations (Supplementary File 2). |
| BMI1 | NFE | Studies have shown BMI1 to have an important function as a biomarker of cancer stem cells with its nuclear expression in laryngeal carcinoma showing correlations with lymph node metastasis. Laryngeal tumors have been found to be common in southern Europe. (64) |
| CISD1 | NFE | Screening of serum prostate-specific antigen levels help in early diagnosis and improved management of prostate cancer. CISD1 was found to be involved in influencing serum PSA levels specifically in European subpopulation. (65) |
| CSNK2A2 | SAS | Telomere length is a heritable trait. Short telomere length has been associated with cardiometabolic risks leading to multiple chronic diseases like Type 2 Diabetes (T2D) which has high incidence rates in South Asians. Recent studies have found CSNK2A2 to be associated with relative leukocyte telomere length in South Asians. (66,67) |
| RNF2 | SAS | Esophageal carcinoma is one of the most frequent malignant tumors and has been found to have high incidence rates in Asia. Studies have found RNF2 to be overexpressed in esophageal carcinoma tissues (68). This gene has a single mutation (T297P) with an allele frequency of 0.01 in FIN subpopulation (Supplementary File 2). |
| SPINK7 | EAS | It is known that East Asian skin is highly sensitive and more reactive than that of Americans and Africans. The weak skin barrier strength and low degree of maturation of the East Asian skin are responsible for increased sensitivity and increased susceptibility to skin inflammations. Therefore, it is obvious that East Asians would have evolved mechanisms to resist skin inflammations. One such prospective candidate that might be involved in resisting inflammations is SPINK7, a serine protease inhibitor and is specific to the East Asian population. This gene was earlier thought to be a tumour suppressor gene, but recent studies show its role in skin homeostasis and inflammatory skin diseases. There is an upregulation of SPINK7 gene expression in inflammatory skin diseases and this may play a role in counterbalancing proteolysis that occurs in these diseases. Thus, the specificity of SPINK7 gene to the normal East Asian population can be associated to resist skin inflammatory diseases which are more prevalent in this population (69,70). This gene has a single mutation (C11R) with an allele frequency of 0.03 in AFR subpopulation (Supplementary File 2). |
| IFITM | EAS | Asia has always been the cradle of influenza right from the historic times. Among the various parts of Asia, Eastern and Southeastern Asia are the main sources of new subtypes of influenza and seasonal influenza that eventually spread to the rest of the world resulting in epidemics/pandemics. One of the resistance mechanisms to combat influenza involves IFITM gene which is specific to this sub-population. IFITM gene codes for a family of interferon- induced antiviral proteins. These transmembrane proteins restrict the viral entry into the cell by preventing them from traversing the cell's lipid bilayer. Also, several SNPs in the IFITM gene are associated with severity in influenza infection and some play an important role in viral restriction. Thus, the specific expression of IFITM gene in East Asians serves as a tool to battle influenza. (71–73) |
| DEFB | EAS | Beta-defensins are a class of antimicrobial proteins that direct the cross-talk between microbes and host. They are produced by epithelial and immune cells. These proteins are encoded by a set of beta-defensin genes (DEFB) that are clustered together in a copy number variable (CNV) as a block. A recent study shows that variation in rapidly evolving sequences of a particular CNV block results in an increased expression of hbd-3 protein. This protein forms a protective surface barrier on epithelial cells and prevents fusion of influenza virus with the host cell. Interestingly, East Asians show a high frequency of DEFB expressing copies as a result of the variation in CNV. This could probably be as a result of selection for resistance to influenza as this region is more prone to influenza and hence, this gene is specific to East Asians. (74,75) |

## DISCUSSION

The ExAC database is an exome sequencing dataset aggregated from large-scale sequencing projects for various population genetic studies such as the Myocardial Infarction Genetics Consortium, Swedish Schizophrenia and Bipolar Studies, NHLBI-GO Exome Sequencing Project, T2D-GENES, SIGMA-T2D and The Cancer Genome Atlas. The dataset includes normal tissue samples (tumor or severe pediatric disease samples were excluded) while samples from late onset diseases such as Type 2 Diabetes and Schizophrenia were included as studies have shown that in these complex diseases, environment plays a major role than genetic factors (76, 77).

The allele frequency spectrum of variations in the dataset revealed rare variations in large excess (Fig. 1), agreeing with the proposition that the age of majority of rare variants in human population is considerably less than that of the common alleles resulting due to an explosive human population growth around 5,000 years ago (78). Of the 10 million variations present in the dataset, variations in each subpopulation having an allele frequency of 0.0001 or greater were used for analysis. Of these, all the non-synonymous variations above the threshold were eliminated to identify invariant genes with no amino acid change at the protein level for each subpopulation.

The analysis revealed a total union set of 1,336 genes across 6 (AFR, AMR, FIN, NFE, SAS, EAS) subpopulations. The inbred FIN subpopulation had 857 invariant genes due to a reduced genetic variation spectrum in its bottlenecked subpopulation (79). Our analysis of the common variants of Finnish invariant genes in other subpopulations showed the role of environmental factors leading to natural selection among these genes. Even though AFR, EAS and SAS represented a diverse population background, we could account for around 250 invariant genes in these subpopulations. Thereby, the variance in these genes in subpopulations did not reflect the population size but rather the extent of diversity present among them.

However, it is interesting to note that each of the subpopulations had certain unique invariant gene sets, some of whose association with diseases have been provided in Table 3. Any variation in genes we have identified as essential in specific subpopulations would result in high risk of diseases uncovered in the ExAC database. Some of the invariant genes arising from our analysis must have withstood the selection pressure and could be essential. These invariant genes are seen to be involved in fundamental processes of the genome functions. Among the molecular functions, we found the involvement of these genes in transcription and translation regulation.

Even though the average exon size of the invariant genes were similar, the average number of exons per invariant gene were found to be much less than the global average. Proteins having large number of exons through alternate splicing exhibit large subtypes and are seen to have diversified functions (80). On the other hand, smaller number of exons as seen in highly evolved proteins such as haemoglobin, myoglobin, alpha globin, insulin etc. Thereby, we believe that the reported invariant genes with lesser number of exons are indeed evolutionarily mature and thus are essential tools in performing functionally crucial fundamental processes.

A total of 46 of 423 invariant genes were found to be absolutely invariant across all subpopulations with a majority from the X chromosome which might be explained by the genes being under a higher selection pressure due to X-linked recessive variants being exposed in males (81, 82). These 46 invariant genes broadly code for two categories of proteins, proteins which are small and compact while the other class of proteins are pretty large (500 to 1,138 amino acids). Most often, the small proteins are generally transcription factors or are those which are present in the Y chromosomes and expressed in the testes (Fig. 4c). In general, the gene sequences coding for these proteins are evolutionarily stable, act as housekeeping genes which are generally expressed in all living cells. The large proteins, on the other hand, are tissue-specific and are essential for the normal functioning of the respective tissues (Fig. 4c). However, even though these proteins vary in sizes, they do not accumulate any mutation. This suggests that the structural plasticity of these proteins is extremely low and any non-synonymous mutation in the gene may lead to a functional loss. X-Ray crystal structures of these proteins might help to reveal more insights into the reason behind their low tolerance for mutations. However, the complete X-Ray crystal structures for these proteins were unavailable thereby the secondary and tertiary structures of these proteins couldn’t be studied.

To our surprise we notice that gene sets of CT47A (Cancer/Testis Antigen Family 47), USP17L (Ubiquitin Specific Peptidase 17), GAGE (G antigen) and PPIAL4 (Peptidylprolyl isomerase) showcased zero expression levels in the GTEx database. Mouse embryonic gene expression databases also showed no entries for these genes. The functional role of these genes set in the evolutionary context remains to be explored. Although these genes are invariant across the subpopulations, their lack of expression may either be due to the fact that these genes must have done their role in the early stages of development (fertilization, spermatogenesis, cell position, morphology etc.) and are no longer required, or their expression needs specific conditions or environmental challenge (deubiquitination, transcription factors etc.). If any mutations occur in these essential genes, it might end up being lethal and thus won’t be seen in the population. In the other case, there hasn’t been a necessary external stimulus needed to provoke the gene’s expression. By this we mean that the gene has either been highly optimized for its function or its expression has not yet been captured.

In conclusion, these 423 invariant genes comprising 2094 exons which are critical in genome function are also likely to be hotspots and any mutation on these would have likely implications in various disease processes including development, cell growth and loss of regulation. Further, of the 46 absolute invariant genes that we identified in 6 subpopulations, 8 were unique to human genome wherein 3 of them still had unknown functions. Although we don’t have any cancer genome representation in the dataset, we notice many of the invariant genes to show association with several cancers and dominant genetic disorders. This set of genes can be used for universal screening for risk for developmental diseases and cancer.

## ACKNOWLEDGMENTS

We thank Sridhar Sivasubbu, Vinod Scaria and Debasis Dash for their valuable inputs in the project. We would also like to thank Vani Brahmachari for her intellectual support and critical reviews of the manuscript.

## CONTRIBUTIONS

SKB conceptualized, designed and guided the project. AKP did the data analysis and AKJ interpreted the results. Manuscript was written and approved by all the authors.

## REFERENCES

1. Gartler SM. The chromosome number in humans: a brief history. Nat Rev Genet. 2006 Aug;7(8):655–60.

2. Seemungal D, Newton G. Human Genome Project. In: Brenner S, Miller JH, editors. Encyclopedia of Genetics [Internet]. New York: Academic Press; 2001 [cited 2019 Aug 17]. p. 980–1. Available from: http://www.sciencedirect.com/science/article/pii/B0122270800017432

3. The 1000 Genomes Project Consortium. A map of human genome variation from population-scale sequencing. Nature. 2010 Oct;467(7319):1061–73.

4. Frazer KA, Murray SS, Schork NJ, Topol EJ. Human genetic variation and its contribution to complex traits. Nat Rev Genet. 2009 Apr;10(4):241–51.

5. Spielman RS, Bastone LA, Burdick JT, Morley M, Ewens WJ, Cheung VG. Common genetic variants account for differences in gene expression among ethnic groups. Nat Genet. 2007 Feb;39(2):226–31.

6. Gymrek M, Willems T, Guilmatre A, Zeng H, Markus B, Georgiev S, et al. Abundant contribution of short tandem repeats to gene expression variation in humans. Nat Genet. 2016 Jan;48(1):22–9.

7. Guttman M, Amit I, Garber M, French C, Lin MF, Feldser D, et al. Chromatin signature reveals over a thousand highly conserved large non-coding RNAs in mammals. Nature. 2009 Mar;458(7235):223–7.

8. Benko S, Fantes JA, Amiel J, Kleinjan D-J, Thomas S, Ramsay J, et al. Highly conserved non-coding elements on either side of *SOX9* associated with Pierre Robin sequence. Nat Genet. 2009 Mar;41(3):359–64.

9. Woolfe A, Goodson M, Goode DK, Snell P, McEwen GK, Vavouri T, et al. Highly Conserved Non-Coding Sequences Are Associated with Vertebrate Development. PLOS Biol. 2004 Nov 11;3(1):e7.

10. Taft RJ, Pang KC, Mercer TR, Dinger M, Mattick JS. Non-coding RNAs: regulators of disease. J Pathol. 2010;220(2):126–39.

11. Sinha P, Jaiswal P, Jainarayanan AK, Brahmachari SK. Intronic miRNA mediated gene expression regulation controls protein crowding inside the cell. Gene. 2018 Dec 30;679:172–8.

12. Lek M, Karczewski KJ, Minikel EV, Samocha KE, Banks E, Fennell T, et al. Analysis of protein-coding genetic variation in 60,706 humans. Nature. 2016 Aug;536(7616):285–91.

13. Casper J, Zweig AS, Villarreal C, Tyner C, Speir ML, Rosenbloom KR, et al. The UCSC Genome Browser database: 2018 update. Nucleic Acids Res. 2018 Jan 4;46(D1):D762–9.

14. Mi H, Muruganujan A, Casagrande JT, Thomas PD. Large-scale gene function analysis with the PANTHER classification system. Nat Protoc. 2013 Aug;8(8):1551–66.

15. Lonsdale J, Thomas J, Salvatore M, Phillips R, Lo E, Shad S, et al. The Genotype-Tissue Expression (GTEx) project. Nat Genet. 2013 May 29;45:580–5.

16. Wang K, Li M, Hakonarson H. ANNOVAR: functional annotation of genetic variants from high-throughput sequencing data. Nucleic Acids Res. 2010 Sep 1;38(16):e164–e164.

17. Bomba L, Walter K, Soranzo N. The impact of rare and low-frequency genetic variants in common disease. Genome Biol. 2017 Apr 27;18(1):77.

18. Kim D, Pertea G, Trapnell C, Pimentel H, Kelley R, Salzberg SL. TopHat2: accurate alignment of transcriptomes in the presence of insertions, deletions and gene fusions. Genome Biol. 2013 Apr 25;14(4):R36.

19. Christiansen JH, Yang Y, Venkataraman S, Richardson L, Stevenson P, Burton N, et al. EMAGE: a spatial database of gene expression patterns during mouse embryo development. Nucleic Acids Res. 2006 Jan 1;34(suppl_1):D637–41.

20. Strikoudis A, Lazaris C, Trimarchi T, Galvao Neto AL, Yang Y, Ntziachristos P, et al. Regulation of transcriptional elongation in pluripotency and cell differentiation by the PHD-finger protein Phf5a. Nat Cell Biol. 2016 Nov;18(11):1127–38.

21. Trappe R, Schulze E, Rzymski T, Fröde S, Engel W. The Caenorhabditis elegans ortholog of human PHF5a shows a muscle-specific expression domain and is essential for C. elegans morphogenetic development. Biochem Biophys Res Commun. 2002 Oct 4;297(4):1049–57.

22. Wang C-Y, Shahi P, Huang JTW, Phan NN, Sun Z, Lin Y-C, et al. Systematic analysis of the achaete-scute complex-like gene signature in clinical cancer patients. Mol Clin Oncol. 2017 Jan 1;6(1):7–18.

23. Guillemot F, Lo L-C, Johnson JE, Auerbach A, Anderson DJ, Joyner AL. Mammalian achaete-scute homolog 1 is required for the early development of olfactory and autonomic neurons. Cell. 1993 Nov 5;75(3):463–76.

24. Mizuguchi R, Kriks S, Cordes R, Gossler A, Ma Q, Goulding M. Ascl1 and Gsh1/2 control inhibitory and excitatory cell fate in spinal sensory interneurons. Nat Neurosci. 2006 Jun;9(6):770–8.

25. Martinelli VC, Kyle WB, Kojic S, Vitulo N, Li Z, Belgrano A, et al. ZASP Interacts with the Mechanosensing Protein Ankrd2 and p53 in the Signalling Network of Striated Muscle. PLOS ONE. 2014 Mar 19;9(3):e92259.

26. Pünzeler S, Link S, Wagner G, Keilhauer EC, Kronbeck N, Spitzer RM, et al. Multivalent binding of PWWP2A to H2A.Z regulates mitosis and neural crest differentiation. EMBO J. 2017 Aug 1;36(15):2263–79.

27. Burrows JF, McGrattan MJ, Johnston JA. The DUB/USP17 deubiquitinating enzymes, a multigene family within a tandemly repeated sequence. Genomics. 2005 Apr 1;85(4):524–9.

28. Chen R, Zhang L, Zhong B, Tan B, Liu Y, Shu H-B. The ubiquitin-specific protease 17 is involved in virus-triggered type I IFN signaling. Cell Res. 2010 Jul;20(7):802–11.

29. Shang B, Gao A, Pan Y, Zhang G, Tu J, Zhou Y, et al. CT45A1 acts as a new proto-oncogene to trigger tumorigenesis and cancer metastasis. Cell Death Dis. 2014 Jun;5(6):e1285.

30. Tang F, Tang S, Guo X, Yang C, Jia K. CT45A1 siRNA silencing suppresses the proliferation, metastasis and invasion of lung cancer cells by downregulating the ERK/CREB signaling pathway. Mol Med Rep. 2017 Nov 1;16(5):6708–14.

31. Chen YT, Iseli C, Yenditti CA, Old LJ, Simpson AJG, Jongeneel C. Identification of a new cancer/testis gene family, CT47, among expressed multicopy genes on the human X chromosome. Genes Chromosomes Cancer. 2006 Apr 1;45(4):392–400.

32. Zhang Q, Su B. Evolutionary Origin and Human-Specific Expansion of a Cancer/Testis Antigen Gene Family. Mol Biol Evol. 2014 Sep 1;31(9):2365–75.

33. Gjerstorff MF, Ditzel HJ. An overview of the GAGE cancer/testis antigen family with the inclusion of newly identified members. Tissue Antigens. 2008 Mar;71(3):187–92.

34. Dave B, Gonzalez DD, Liu Z-B, Li X, Wong H, Granados S, et al. Role of RPL39 in Metaplastic Breast Cancer. J Natl Cancer Inst. 2017 Jun;109(6):djw292.

35. Desruisseaux MS, Iacobas DA, Iacobas S, Mukherjee S, Weiss LM, Tanowitz HB, et al. Alterations in the Brain Transcriptome in Plasmodium Berghei ANKA Infected Mice. J Neuroparasitology. 2010 Oct;1.

36. Dkhil MA, Al-Shaebi EM, Lubbad MY, Al-Quraishy S. Impact of sex differences in brain response to infection with Plasmodium berghei. Parasitol Res. 2016 Jan 1;115(1):415–22.

37. Yamashita Y, Nishiumi S, Kono S, Takao S, Azuma T, Yoshida M. Differences in elongation of very long chain fatty acids and fatty acid metabolism between triple-negative and hormone receptor-positive breast cancer. BMC Cancer. 2017 Aug 29;17(1):589.

38. Gong J, Schumacher F, Lim U, Hindorff LA, Haessler J, Buyske S, et al. Fine Mapping and Identification of BMI Loci in African Americans. Am J Hum Genet. 2013 Oct 3;93(4):661–71.

39. Pillay V, Crowther NJ, Ramsay M, Smith GD, Norris SA, Lombard Z. Exploring genetic markers of adult obesity risk in black adolescent South Africans—the Birth to Twenty Cohort. Nutr Diabetes. 2015 Jun;5(6):e157.

40. Graff M, North KE, Richardson AS, Young KM, Mohlke KL, Lange LA, et al. Screen time behaviours may interact with obesity genes, independent of physical activity, to influence adolescent BMI in an ethnically diverse cohort. Pediatr Obes. 2013;8(6):e74–9.

41. Udpa N, Ronen R, Zhou D, Liang J, Stobdan T, Appenzeller O, et al. Whole genome sequencing of Ethiopian highlanders reveals conserved hypoxia tolerance genes. Genome Biol. 2014 Feb 20;15(2):R36.

42. Manku H, Langefeld CD, Guerra SG, Malik TH, Alarcon-Riquelme M, Anaya J-M, et al. Trans-Ancestral Studies Fine Map the SLE-Susceptibility Locus TNFSF4. PLOS Genet. 2013 Jul 18;9(7):e1003554.

43. Sanchez E, Comeau ME, Freedman BI, Kelly JA, Kaufman KM, Langefeld CD, et al. Identification of novel genetic susceptibility loci in African-American lupus patients using a candidate gene association study. Arthritis Rheum. 2011 Nov;63(11):3493–501.

44. Hinterseher I, Erdman R, Elmore JR, Stahl E, Pahl MC, Derr K, et al. Novel pathways in the pathobiology of human abdominal aortic aneurysms. Pathobiol J Immunopathol Mol Cell Biol. 2013;80(1):1–10.

45. Li H, Li J, Su Y, Fan Y, Guo X, Li L, et al. A novel 3p22.3 gene *CMTM7* represses oncogenic EGFR signaling and inhibits cancer cell growth. Oncogene. 2014 Jun;33(24):3109–18.

46. Verma M. Epigenome-Wide Association Studies (EWAS) in Cancer. Curr Genomics. 2012 Jun;13(4):308–13.

47. Hulur I, Gamazon ER, Skol AD, Xicola RM, Llor X, Onel K, et al. Enrichment of inflammatory bowel disease and colorectal cancer risk variants in colon expression quantitative trait loci. BMC Genomics. 2015 Feb 27;16:138.

48. Peltekova VD, Lemire M, Qazi AM, Zaidi SHE, Trinh QM, Bielecki R, et al. Identification of genes expressed by immune cells of the colon that are regulated by colorectal cancer-associated variants. Int J Cancer. 2014 May 15;134(10):2330–41.

49. Nair KS, Bigelow ML, Asmann YW, Chow LS, Coenen-Schimke JM, Klaus KA, et al. Asian Indians Have Enhanced Skeletal Muscle Mitochondrial Capacity to Produce ATP in Association With Severe Insulin Resistance. Diabetes. 2008 May 1;57(5):1166–75.

50. Sapienza C, Lee J, Powell J, Erinle O, Yafai F, Reichert J, et al. DNA methylation profiling identifies epigenetic differences between diabetes patients with ESRD and diabetes patients without nephropathy. Epigenetics. 2011 Jan 1;6(1):20–8.

51. Ali HEA, Lung P-Y, Sholl AB, Gad SA, Bustamante JJ, Ali HI, et al. Dysregulated gene expression predicts tumor aggressiveness in African-American prostate cancer patients. Sci Rep. 2018 Nov 5;8(1):1–12.

52. Costa GNO, Dudbridge F, Fiaccone RL, da Silva TM, Conceição JS, Strina A, et al. A genome-wide association study of asthma symptoms in Latin American children. BMC Genet. 2015 Dec 3;16:141.

53. Stechman MJ, Loh NY, Thakker RV. Genetic causes of hypercalciuric nephrolithiasis. Pediatr Nephrol Berl Ger. 2009 Dec;24(12):2321–32.

54. Romero V, Akpinar H, Assimos DG. Kidney stones: a global picture of prevalence, incidence, and associated risk factors. Rev Urol. 2010;12(2–3):e86–96.

55. Anvari S, Vyhlidal CA, Dai H, Jones BL. Genetic Variation along the Histamine Pathway in Children with Allergic versus Nonallergic Asthma. Am J Respir Cell Mol Biol. 2015 Apr 24;53(6):802–9.

56. Elmansy D, Koyutürk M. Cross-population analysis for functional characterization of type II diabetes variants. BMC Bioinformatics. 2019 Jun 20;20(12):320.

57. Marttila M, Kytövuori L, Helisalmi S, Kallio M, Laitinen M, Hiltunen M, et al. Molecular Epidemiology of Charcot-Marie-Tooth Disease in Northern Ostrobothnia, Finland: A Population-Based Study. Neuroepidemiology. 2017;49(1–2):34–9.

58. Särkijärvi S, Kuusisto H, Paalavuo R, Levula M, Airla N, Lehtimäki T, et al. Gene expression profiles in Finnish twins with multiple sclerosis. BMC Med Genet. 2006 Feb 27;7(1):11.

59. Ryu EJ, Wang JYT, Le N, Baloh RH, Gustin JA, Schmidt RE, et al. Misexpression of Pou3f1 Results in Peripheral Nerve Hypomyelination and Axonal Loss. J Neurosci. 2007 Oct 24;27(43):11552–9.

60. Vavougios GD, Zarogiannis SG, Krogfelt KA, Gourgoulianis K, Mitsikostas DD, Hadjigeorgiou G. Novel candidate genes of the PARK7 interactome as mediators of apoptosis and acetylation in multiple sclerosis: An in silico analysis. Mult Scler Relat Disord. 2018 Jan 1;19:8–14.

61. Häkli S, Luotonen M, Sorri M, Majamaa K. Mutations in the two ribosomal RNA genes in mitochondrial DNA among Finnish children with hearing impairment. BMC Med Genet. 2015 Feb 4;16:3.

62. Gerits A, Nieminen P, Muynck SD, Carels C. Exclusion of coding region mutations in MSX1, PAX9 and AXIN2 in eight patients with severe oligodontia phenotype. Orthod Craniofac Res. 2006;9(3):129–36.

63. Nieminen P, Arte S, Pirinen S, Peltonen L, Thesleff I. Gene defect in hypodontia: exclusion of MSX1 and MSX2 as candidate genes. Hum Genet. 1995 Sep 1;96(3):305–8.

64. Allegra E, Trapasso S. Nuclear BMI1 as a Biomarker in Laryngeal Cancer. In: Preedy VR, Patel VB, editors. Biomarkers in Cancer [Internet]. Dordrecht: Springer Netherlands; 2014 [cited 2019 Aug 16]. p. 1–8. Available from: https://doi.org/10.1007/978-94-007-7744-6_15-1

65. Ge Y-Z, Xu Z, Xu L-W, Yu P, Zhao Y, Xin H, et al. Pathway analysis of genome-wide association study on serum prostate-specific antigen levels. Gene. 2014 Nov 1;551(1):86–91.

66. Sethi I, Bhat GR, Singh V, Kumar R, Bhanwer AJS, Bamezai RNK, et al. Role of telomeres and associated maintenance genes in Type 2 Diabetes Mellitus: A review. Diabetes Res Clin Pract. 2016 Dec 1;122:92–100.

67. Saxena Richa, Bjonnes Andrew, Prescott Jennifer, Dib Patrick, Natt Praveen, Lane Jacqueline, et al. Genome-Wide Association Study Identifies Variants in Casein Kinase II (CSNK2A2) to be Associated With Leukocyte Telomere Length in a Punjabi Sikh Diabetic Cohort. Circ Cardiovasc Genet. 2014 Jun 1;7(3):287–95.

68. Yang X-X, Ma M, Sang M-X, Wang X-X, Song H, Liu Z-K, et al. Radiosensitization of esophageal carcinoma cells by knockdown of RNF2 expression. Int J Oncol. 2016 May 1;48(5):1985–96.

69. Weber C, Fischer J, Redelfs L, Rademacher F, Harder J, Weidinger S, et al. The serine protease inhibitor of Kazal-type 7 (SPINK7) is expressed in human skin. Arch Dermatol Res. 2017 Nov 1;309(9):767–71.

70. Muizzuddin N, Hellemans L, Van Overloop L, Corstjens H, Declercq L, Maes D. Structural and functional differences in barrier properties of African American, Caucasian and East Asian skin. J Dermatol Sci. 2010 Aug 1;59(2):123–8.

71. Wen F, Bedford T, Cobey S. Explaining the geographical origins of seasonal influenza A (H3N2). Proc Biol Sci. 2016 14;283(1838).

72. Bailey CC, Zhong G, Huang I-C, Farzan M. IFITM-Family Proteins: The Cell’s First Line of Antiviral Defense. Annu Rev Virol. 2014 Nov 1;1:261–83.

73. Wellington D, Laurenson-Schafer H, Abdel-Haq A, Dong T. IFITM3: How genetics influence influenza infection demographically. Biomed J. 2019 Feb 1;42(1):19–26.

74. Meade KG, O’Farrelly C. β-Defensins: Farming the Microbiome for Homeostasis and Health. Front Immunol [Internet]. 2019 [cited 2019 Aug 12];9. Available from: https://www.frontiersin.org/articles/10.3389/fimmu.2018.03072/full

75. Hardwick RJ, Machado LR, Zuccherato LW, Antolinos S, Xue Y, Shawa N, et al. A worldwide analysis of beta-defensin copy number variation suggests recent selection of a high-expressing DEFB103 gene copy in East Asia. Hum Mutat. 2011 Jul;32(7):743–50.

76. Dendup T, Feng X, Clingan S, Astell-Burt T. Environmental Risk Factors for Developing Type 2 Diabetes Mellitus: A Systematic Review. Int J Environ Res Public Health. 2018 Jan;15(1):78.

77. Morgan C, Fisher H. Environment and Schizophrenia: Environmental Factors in Schizophrenia: Childhood Trauma—A Critical Review. Schizophr Bull. 2007 Jan 1;33(1):3–10.

78. Fu W, Jun G, Kang HM, Abecasis G, Leal SM, Gabriel S, et al. Analysis of 6,515 exomes reveals the recent origin of most human protein-coding variants. Nature. 2013 Jan;493(7431):216–20.

79. Lim ET, Würtz P, Havulinna AS, Palta P, Tukiainen T, Rehnström K, et al. Distribution and Medical Impact of Loss-of-Function Variants in the Finnish Founder Population. PLOS Genet. 2014 Jul 31;10(7):e1004494.

80. Romero PR, Zaidi S, Fang YY, Uversky VN, Radivojac P, Oldfield CJ, et al. Alternative splicing in concert with protein intrinsic disorder enables increased functional diversity in multicellular organisms. Proc Natl Acad Sci U S A. 2006 May 30;103(22):8390–5.

81. Johnson NA, Lachance J. The genetics of sex chromosomes: evolution and implications for hybrid incompatibility. Ann N Y Acad Sci. 2012 May;1256:E1–22.

82. Veeramah KR, Gutenkunst RN, Woerner AE, Watkins JC, Hammer MF. Evidence for Increased Levels of Positive and Negative Selection on the X Chromosome versus Autosomes in Humans. Mol Biol Evol. 2014 Sep;31(9):2267–82.

