## Supplementary Figure 1 for "Invariant Genes in Human Genomes"

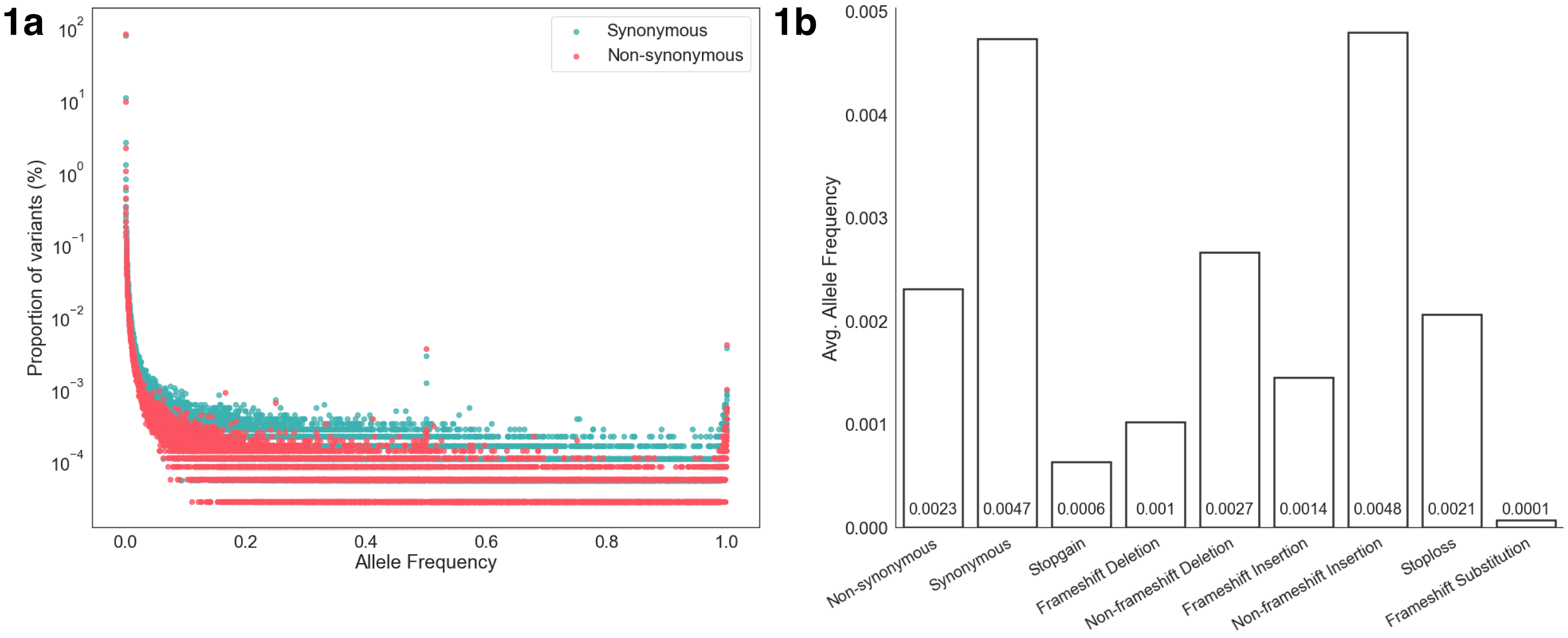


**Supplementary Figure 1 I a,** Distribution of synonymous and non-synonymous variations in ExAC dataset with respect to their allele frequencies rounded off to 4 decimal places. **b,** Average allele frequency of coding mutations in the ExAC dataset.
