## Supplementary Figure 2 for "Invariant Genes in Human Genomes"

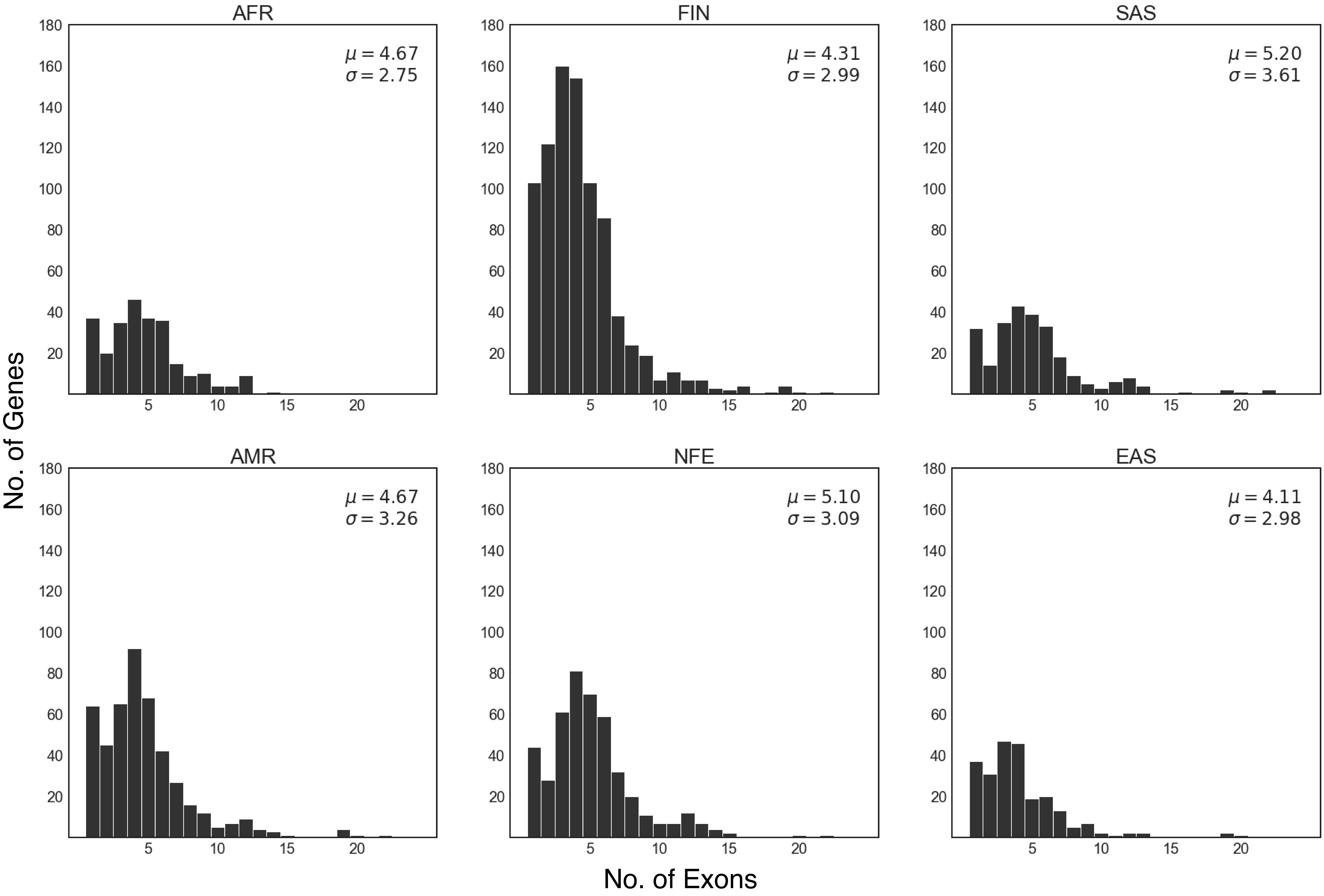


**Supplementary Figure 2 |** Distribution of number of exons per gene in invariant genes across subpopulations.
