## Supplementary Figure 3 for "Invariant Genes in Human Genomes"

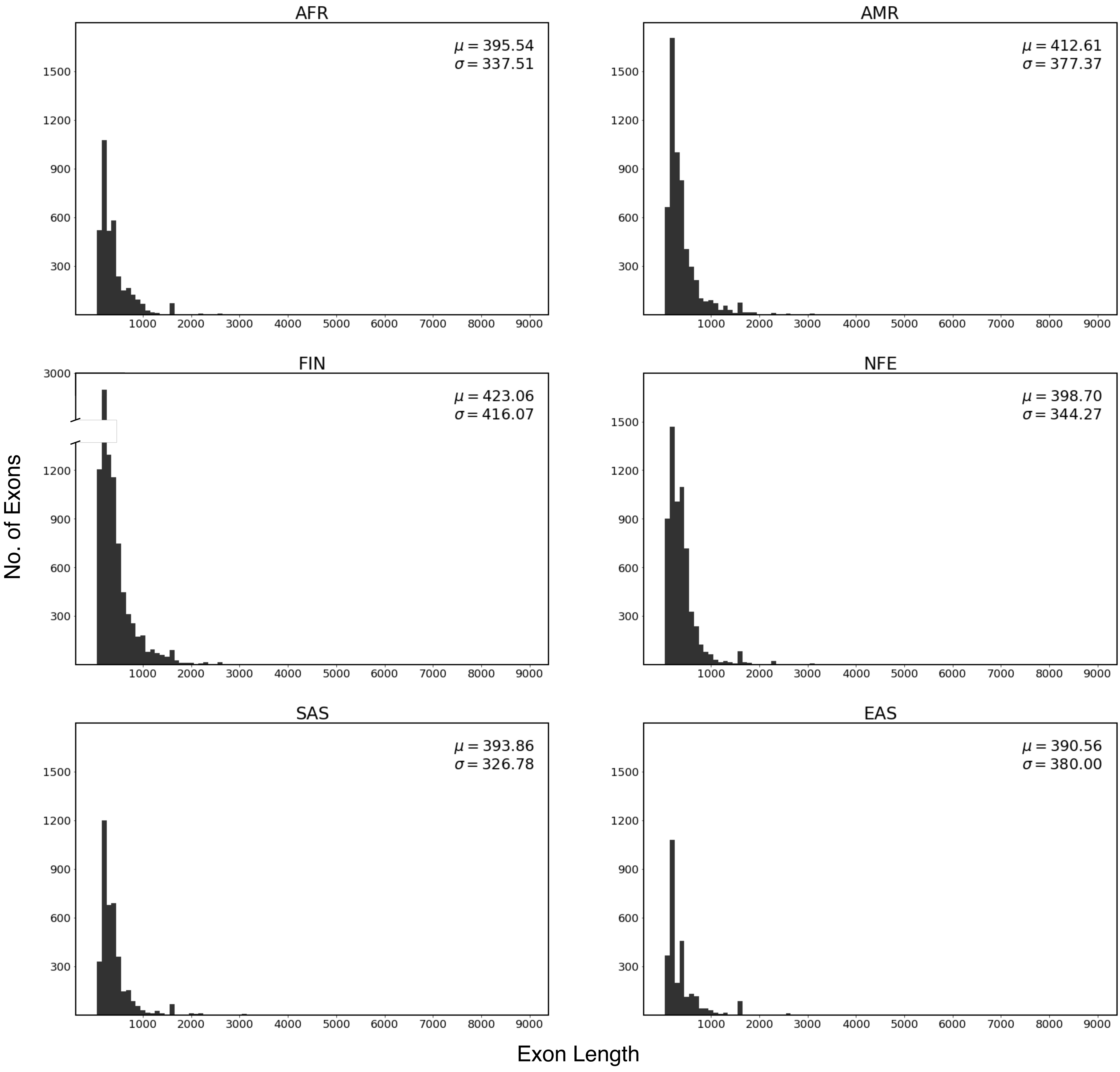


**Supplementary Figure 3 l** Distribution of exon lengths in invariant genes across subpopulations.
