## Supplementary Figure 4 for "Invariant Genes in Human Genomes"

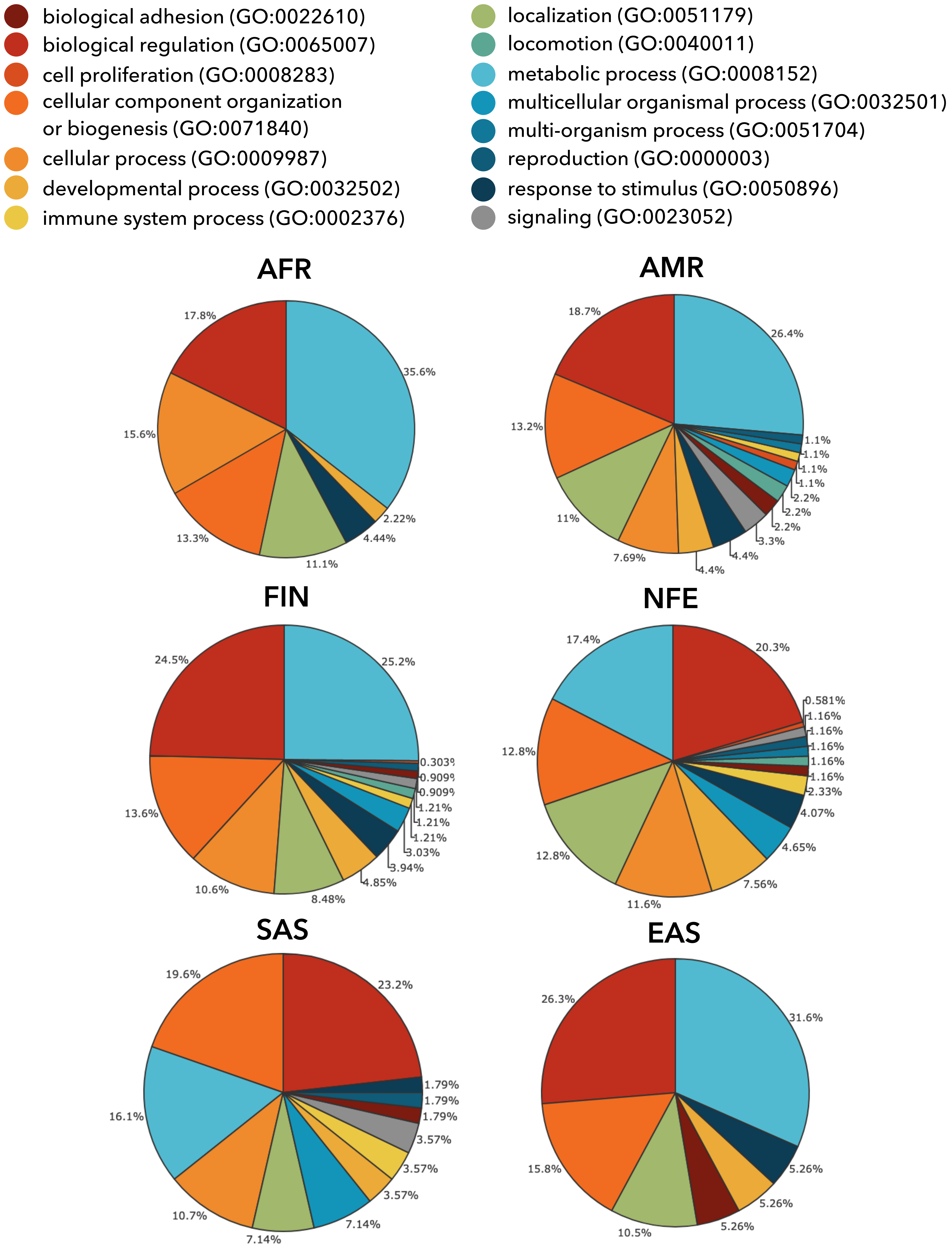


**Supplementary Figure 4 I** Share of biological processes performed by invariant genes specific to each subpopulation.
