## Supplementary Figure 5 for "Invariant Genes in Human Genomes"

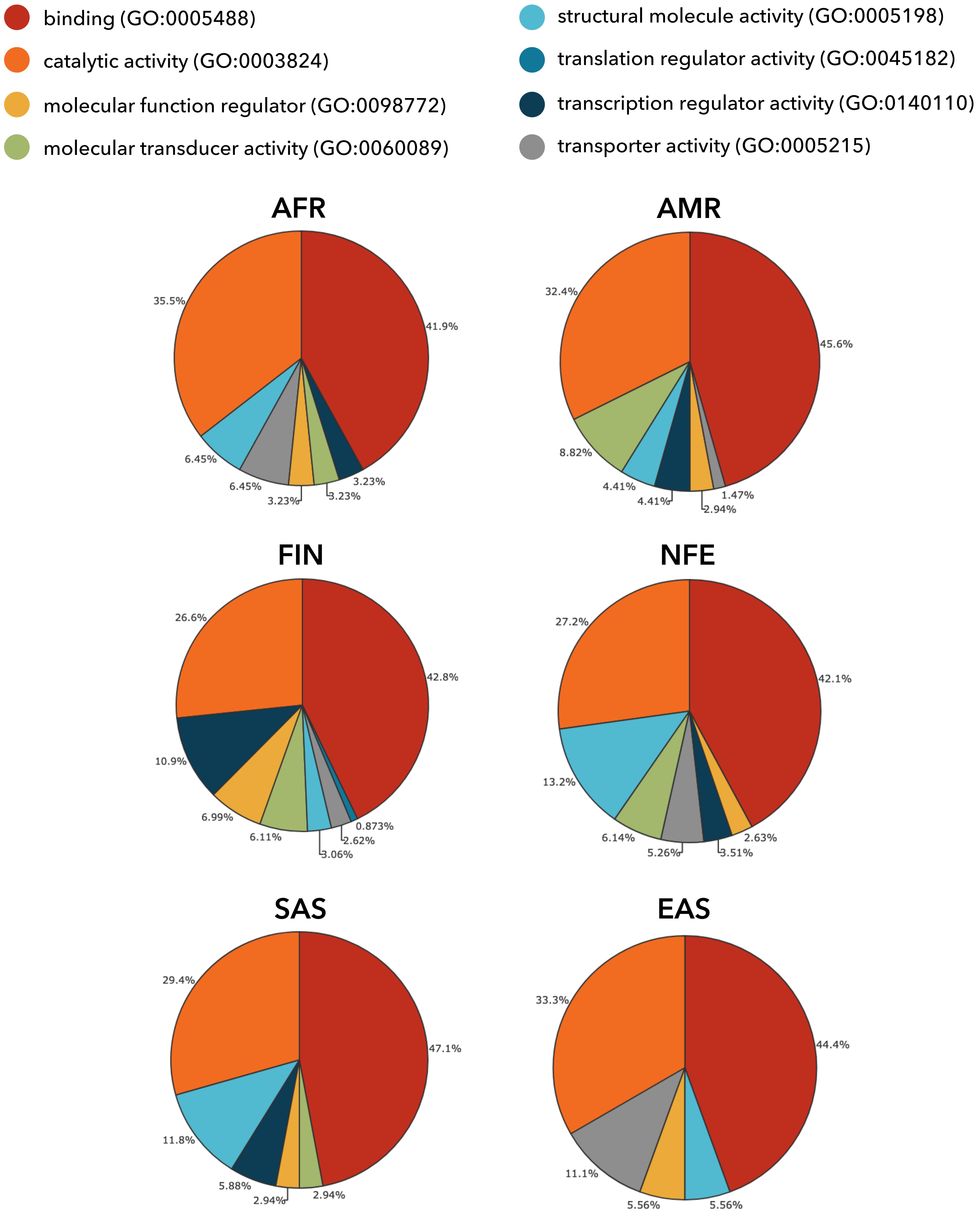


**Supplementary Figure 5 I** Share of molecular functions performed by invariant genes specific to each subpopulation
